# Logistic resource limitation model for quasi real-time measured subjective cognitive load predicts Hill function of hemoglobin-oxygen saturation

**DOI:** 10.1101/2024.01.23.576976

**Authors:** Norbert Fürstenau

**Affiliations:** Institute of Flight Guidance, German Aerospace Center (DLR), Braunschweig, Germany

## Abstract

Cognitive processing and memory resources invested in task execution determine mental workload (MWL) that is quantified through objective physiological measures such as heart rate and variability, EEG, and hemoglobin oxygen (HbO_2_) saturation, and subjective methods like periodic quasi-real-time “instantaneous self-assessment” (ISA) with discrete five- or seven-level WL-scales. Previously published results of human-in-the-loop (HITL) air-traffic control simulations with highly trained domain experts provided initial evidence for logistic and power law functional dependencies between subjective MWL self-assessment reports and simultaneously monitored task load and simulation variables (e.g. communication and traffic load). Here we show that a biased “Logistic Resource Limitation” (LRL) model for regression based parameter estimates of subjective self-reports through combination with a logistic task load function leads to a cognitive power law with parametric correspondence to the classical Hill function that quantifies HbO_2_ saturation. Hill function saturation exponent and equilibrium dissociation constant turned out to show surprising agreement with corresponding estimates of the power law parameters derived from the LRL-model applied to published independent data sets from the three different HITL-simulation experiments. Our results suggest the hypothesis that under certain conditions quasi real-time subjective (behavioral) reporting of cognitive load due to task execution might represent the output of an interoceptive HbO_2_ saturation sensor that measures resource limitation of neural energy supply. From the HbO_2_ - saturation perspective, our results might provide an additional aspect to the “selfish brain” theory for cortical energy supply as derived by A. Peters et al. based on a logistic Glucose push–pull supply chain model. However, more focused experiments are required including direct (e.g. fNIR based) measurements of HbO_2_-saturation to further support (or falsify) our conclusions.

**Author Summary:** Measurements of mental workload of domain experts under cognitive task requirements by human-in-the-loop simulation experiments utilize subjective and objective methods and measures. Standard data analysis is mostly limited to linear statistical methods such as variance and regression analysis for quantifying load differences under different task requirements. Based on nonlinear resource limitation models with asymptotic saturation limits we derive here a cognitive power law for the dependency of real-time subjective work- vs. objective task load. The focused analysis of three previously published independent datasets revealed an unexpected formal and quantitative equivalence with the classical Hill-function of blood-oxygen saturation. Our results suggest the hypothesis of a close quantitative relationship between subjective load reports and an interoceptive senor for cortical energy resources.

## 1. Introduction

In this article we derive a formal parametric equivalence between a cognitive power law of mental load quantified on a subjective behavioral level [1] [2] and the classical Hill function of hemoglobin oxygen (HbO_2_) saturation [3]. Evidence is provided through theoretical parameter predictions and model based data analysis of three previously published independent data sets [1] [2] [4]. Cognitive processing and memory resources (e.g. [5] [6]) invested in task execution determine mental workload (MWL) that is quantified through objective measures such as EEG [7] [8], heart rate variability [8] [9] and HbO_2_ saturation [10], and subjective methods like periodic quasi real-time “instantaneous self-assessment” (ISA) techniques (realizing Kahneman’s “Thinking fast” [11]) with discrete five- or seven level MWL-scales [12] [13] [14]. For quantitatively relating subjective to objective measures formal psycho-physical [1] [15] [16] [17] [18] [19] [20] and (neuro-) physiological models [21] [22] [23] are required to enable theoretical predictions and regression based parameter estimates. The latter authors referred to Friston’s proposal for a unified brain theory based on the (thermodynamic) free energy principle [24], and they provided evidence for a formal logistic supply chain (push-pull) model of Glucose based energy transport into the brain [25], termed “the selfish brain” [22]. They formalized cerebral energy status control by allocative brain pull and suppression of blood to periphery flux in favor of energy supply of the brain under acute demand due to stress, based on peripheral and cerebral energy signals. E.g. through neuronal monitoring of intracellular energy resource (ATP concentration) by Glucose dependent K-channels on the surface of certain neuron types for controlling neural activity [25].

For estimates of MWL and task load (TL) model parameter values in aeronautics experimental evidence is typically obtained from data of realistic human-in-the-loop (HITL) simulations like flight simulation (e.g. [10] [26]) and air traffic control (ATC) [1] [2] [27] [28] [29] [30] [31] [32]. Highly trained domain experts (air traffic control operators, ATCOs) provided subjective MWL-reports together with simultaneously monitored objective task load and environmental simulation variables (e.g. communication and traffic load) [1] [2] [31] and physiological measures [8] [9] [10]. Subjective MWL-reports were monitored in quasi real-time with fixed reporting time-intervals of typically Δt =2 or 5 min by means of discrete 7-level (air traffic workload input technique ATWIT [12] and 5-level ISA workload measures (Instantaneous Self-Assessment and 1 ≤ ISA ≤ 5 [13] [14] [33] [34]). ATC-simulations with subjective MWL-reporting and measurement of physiological data under well-defined cognitive task requirements had shown discriminability of MWL-levels and (linear) correlations [9] [10] [35] [36]. Initial quantitative parametric dependencies quantified by nonlinear models were published recently [1] [2] [4] [32] [37].

Here we show that a biased “Logistic Resource Limitation” (LRL) model for nonlinear regression analysis of subjective quasi-real-time MWL self-reports as function of environmental (traffic) load, through combination with a logistic task load characteristic (objective TL, e.g, rate of radio calls RC) after normalization S(TL) leads to a cognitive power law MWL(S) (see [1] [32] for initial validation results). It exhibits quantitative parametric correspondence to the classical Hill function of HbO_2_-saturation [3] [38].

Regression-based MWL(S)-parameter estimates of the cognitive power law parameters (exponent and logistic shift factor) from HITL-data obtained with three independent ATC-simulation experiments [1] [2] [4] [37] turned out to be of the same order of magnitude as biochemical Hill-function parameters (HbO_2_-saturation exponent, equilibrium dissociation constant) [3].

This agreement suggests the hypothesis that subjective MWL-reporting might represent the output of an interoceptive HbO_2_ saturation sensor [39] indicating neural energy resource limitation, possibly related to cerebral control mechanisms as described in [23] [25]. Moreover, after MWL-normalization the new subjective response variable R(MWL) as function of objective cognitive stimulus S(TL) represents a cognitive psychophysical Stevens law with the same (Hill) exponent that was derived and validated in [1] [2]. It corresponds to Stevens’ classical (physical) stimulus – (subjective) response power law characteristic [15] [19] [20]. This relationship confirmed the early result of Gopher et al. concerning the psychophysical foundation of cognitive load [16], and it provides a formal link to excitable network based power law functions as microscopic foundation of psychophysical laws [17] [18]. Experimental evidence for cognitive effort with corresponding Glucose based energy consumption to significantly influence decision making was obtained in research on embodiment of cognition [40] [41].

This Introduction is followed by Theory-section 2 with derivation of the cognitive Hill equation and the related Stevens law, based on the LRL-model. Results of logistic and derived Hill-function parameter estimates based on three previously published independent data sets and comparison with the classical HbO_2_ dissociation curve parameters are presented in section 3 and discussed in section 4. A conclusion is provided in section 5. The methods used in the three previously published ATC-simulation experiments for testing the predictions of the LRL-model are reviewed briefly in a supplement that also includes tables with preprocessed data for experiments 2 and 3. For experiment 1 the data are available online in [1].

## 2. Methods: Theory

In the following five subsections we derive the formal and quantitative equivalence between the cognitive power law dependency between subjective mental workload and objective task load (cognitive Hill function) and Hill’s classical physiological equation for hemoglobin-oxygen (HbO_2_) saturation. For this purpose we assume the individual subjective discrete quasi-online MWL-reports (five or seven level ISA (I = 1…5) or ATWIT (W = 1…7)) to represent values of a continuous scale (see Discussion section 4). For the experimental validation (see section 3) of the theoretical predictions of the LRL-model we use averages across (domain expert) subjects and repeated MWL and TL measures as dependent on environmental (ATC- traffic) load. The logistic MWL function in section 2.2 and the cognitive Hill function in 2.4 to be compared with Hill’s classical function in section 2.5 are extended versions of model functions derived in [1] [2], allowing for a logistic MWL-bias Δ > 0.

### 2.1. Logistic function for cognitive resource limitation

The sigmoid or logistic function as formal basis of theoretical LRL-model predictions and regression based WL-data analysis represents the general solution of the classical Verhulst or logistic nonlinear differential equation (DEQ, e.g. [42]).

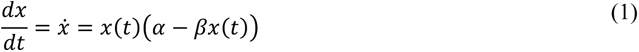

The Verhulst equation may be used as the most simple approach for describing the dynamics of resource limited systems with time t as independent variable (converging to resource limit x_u_ = α/β for lim t→∞). Given α > β, it models a linear differential increase ∼ α x, i.e. exponential growth x ∼ exp{αt} for small t, and when βx converges against α a linear differential decrease ∼ (α – βx), i.e. exponential approach x(t) to the upper asymptote x_u_. The logistic function as general solution (including a time-shift parameter s) is given by

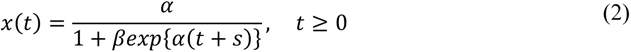

with lower asymptote x_d_ = 0, and initial value (intersection with ordinate) x(t=0) = x_0_ = α/(β + exp{-α s}) < x_u_.

For the cognitive WL-/TL-models within the ATC-simulation context we replace time in x(t) by traffic count or flow n as independent variable (number of aircraft (or AC per time interval AC/Δt entering sector as determined by boundary distance D, speed v, AC-separation distance d_min_, and frequency of handover / clearance requests for entering sector). For simplicity we use n as general independent simulation variable for traffic load. Where necessary we will introduce dimensions and discriminate between flow (AC/Δt) and sector traffic count N_S_, with N_Smax_ = D / d_min_ and n = v/d_min_ = N_S_ v / D (AC/Δt). Typically for the approach sector (Terminal Radar Approach Control, TRACON distance) D = 30 … 50 nautical miles (nm), d_min_ = 3 … 5 nm, v = 120 … 150 nm/h = kn, so that N_Smax_ :≈ 15 and n_c_ :≈ 45 (AC/hour) as (approximate ) capacity limit characterizing a typical high load scenario.

In order for the MWL-sigmoid to allow for the WL-measure intercept I(n = 0) = I_0_ to asymptotically approach (with increasing shift constant s) the lower limit of ATWIT or ISA WL-scales I_d_ := 1 < I_0_, (lower asymptote x_d_ > 0 in equ. (2)) an offset (bias) has to be selected according to lim I(n → - ∞) = Δ ≥ 0 (of course only n ≥ 0 is possible). It results in a WL-scale shift with I_Δ_ = I - Δ. The WL-bias parameter Δ < I_0_ may be assumed to represent a minimum of cognitive tasks, usually present even in the underload range of WL (range of very small n), before the exponential WL(n)-increase becomes noticeable due to increasing task requirements TL(n) of the dominating task variable. Bias Δ is assumed to be sensitive to changes of usually constant work conditions with residual influence on WL even for vanishing effect of the dominating task (as observed in [1]). Including the offset, the Verhulst DEQ formalizing the change of WL with (environmental) independent traffic variable n is given by

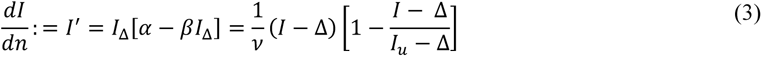

with exponential rate or sensitivity parameter α = 1/ν, saturation parameter β = (ν (I_u_ – Δ))^-1^, and α/β = (I_u_ – Δ). I_u_ defines the (asymptotic) cognitive resource or saturation limit as prior information. For the ISA WL-scale the mental overload range due to unmanageable number of cognitive tasks is defined by I(n) > 4. I ≈ 4 corresponds to a typical traffic capacity limit n_c_ ≈ 45 (AC/h, see above), where also the communication task load (rate of radio calls R(n)) approaches the theoretical limit that may be estimated by considering the average radio call duration of experienced domain experts (≈ 3 – 4 s, see sections 2.3 and 3.1, 3.2 and [43]).

### 2.2. Logistic resource limitation (LRL) model of subjective mental workload

The following equ. (*4*) represents the theoretical ISA-WL characteristic with I_d_ = 1 ≤ I(n) ≤ I_u_ = 5 as solution of the resource limitation DEQ (3):

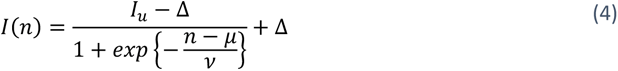

Eq. (*4*) represents the logistic function in terms of the logistic WL-parameters (I_u_, Δ, μ, ν), with constant shift factor k = exp{μ/ν}. In the dimensionless exponent traffic flow n may be exchanged by sector traffic count N_s_. Logistic shift k may be expressed by means of intersection I(n=0) = I_0_ and bias Δ, derived as k = (I_u_ – I_0_)/(I_0_ – Δ). k = 4 for Δ = 0 and I_0_ := I_d_ = 1 by plausibility was selected for the initial validation in [1] (see section 3.1). Figure *1* depicts two representative theoretical sigmoid characteristics with Δ = 0 and 1 respectively.

**Figure 1:**
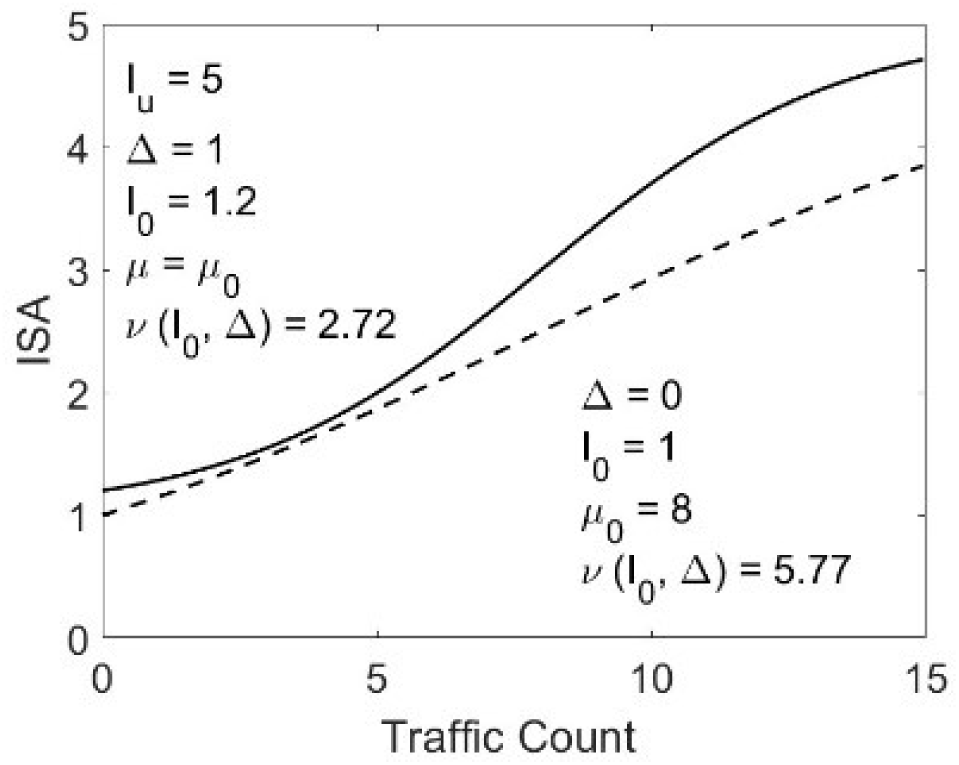
Subjective ISA-MWL report vs. traffic load (theory). Dashed curve: Sigmoid solution of Verhulst equation for MWL ISA-bias Δ = 0 (lower asymptote) and Intersection I_0_ = I_d_ = 1 (lower scale limit). Abscissa scale = traffic load (count N_S_). Solid curve: For Δ = 1 lower asymptote shifts to lower scale limit I_d_ = 1, with intersection I_0_ near Δ. Logistic shift μ_0_ = inversion point near transition to excessive MWL corresponding to unmanageable task load (see Figure 2).

For MWL-bias Δ := 1 < I_0_, k may be a very large number if I_0_ – Δ << 1 or μ >> ν. With I_0_ – Δ << 1 and prior theoretical estimate μ :≈ 7 as example (corresponding to capacity traffic count n_c_ :≈ 7), see following sections 2.3, and 3.2), a theoretical estimate of scaling parameter ν is obtained via ν = μ/ln(k) = 5 for Δ = 0, and ν = 2.4 with k = 19 (for Δ = 1 and I_0_ := 1.2, where we assume a plausible sigmoid intercept range I_0_ = 1. 1 …1.3 for standard work conditions). Δ ≈ 1 was confirmed for the Lee et al. data [4] in section 3.2. With the scaling parameters ν in Figure 1 the sensitivities (slope) dI/dn := I’ of the normalized characteristic I(n)/I_u_ at inversion point N_S_ = μ are obtained as I’ = (I_u_ – Δ)/(4ν) = 0.214 for Δ = 0, and I’ = 1/ν = 0.368 for Δ = 1. For two different previously published HitL-simulations in [1] [2] (experiment 1 and 3 respectively, section 3) we used model equation (*4*) without offset, i.e. Δ := 0 and with intercept I_0_ = 1 as plausible assumption that reduced equ. (*4*) to a regression model with a single unknown parameter to be estimated, due to μ = ν ln(4). For experiment 3 improved parameter estimates with Δ = 1 are provided in section 3.3

### 2.3. LRL-model of radio calls task load

Like equation (*4*) for the subjective MWL-characteristic we can define for the objective TL a logistic characteristic and predict theoretical parameter estimates for ATCo’s rate of radio calls for handovers and clearances (cognitive task load RC = number of calls per Δt, short R(n)) as function of (control sector) traffic count or flow. Radio communication with pilots plays a dominating role in controller’s TL (e.g. [27] [28] [29]; see also [2]). In experiment 1 ( [1], see section 3.1) it was monitored periodically, synchronously with ISA-reporting in 2 min time steps. An upper asymptotic limit R_u_ of frequency (rate) of radio calls (RC (2 min)^-1^) may be derived using average radio call duration of <RCD= ≈ 4-5 s/call as prior information [1] [2] [43], and assuming equal call duration of ATCO and pilot with minimum interval between calls := 1 s we obtain R_u_ = 120 s /(2 x 5 + 1) s = 10…12 (calls (2 min)^-1^) as reasonable estimate. In previous work [1] scenarios with four levels of traffic flow n suggested the plausible model that traffic RC(n) starts with linear growth, i.e. R(n) ∼ n for n << 10 calls per 2 min. Consequently, this required the sigmoid inflexion point of maximum slope at n = μ_R_ := 0, R(n=0) = R_0_ = 0 (i.e. offset Δ_R_ := -R_u_ (k_R_ = exp(μ_R_/ρ) = 1) and exponential approach to R_u_, represented by:

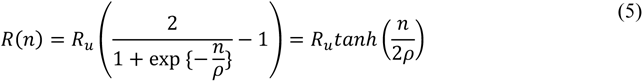

A rough estimate for scaling (TL sensitivity) parameter ρ can be derived from the maximum value of the slope at inversion (dR(n=0)/dn = R’_0_), as *R*_0_^’^ = *^R^_u_*/2ρ≈Δ*R*/Δ*n*. Using tangent slope estimate at R_0_ for extrapolation to intersection with R = R_u_ as: R_0_’ ≈ ΔR(n=0)/n_0_ =R_u_/n_c_, this yields *ρ*: ≈*R_u_*/2*R*_0_’=n_c_/2 = 3.5, and normalized slope Δp_R_/Δn = 1 / 2ρ ≈ 1/ n_c_ ≈ 0.14. Here we used n_c_ := 7 (or ca. 40 AC/h; dimensionless exponent allows to use n for traffic count and flow as well) as upper boundary of sector traffic count where controllers begin to change communication strategy (according to prior information from domain experts, see section 3.2, [1] [30] [37]). The following *Figure 2* depicts the corresponding theoretical prediction for the normalized radio call rate p_R_ := R(n)/R_u_.:

**Figure 2:**
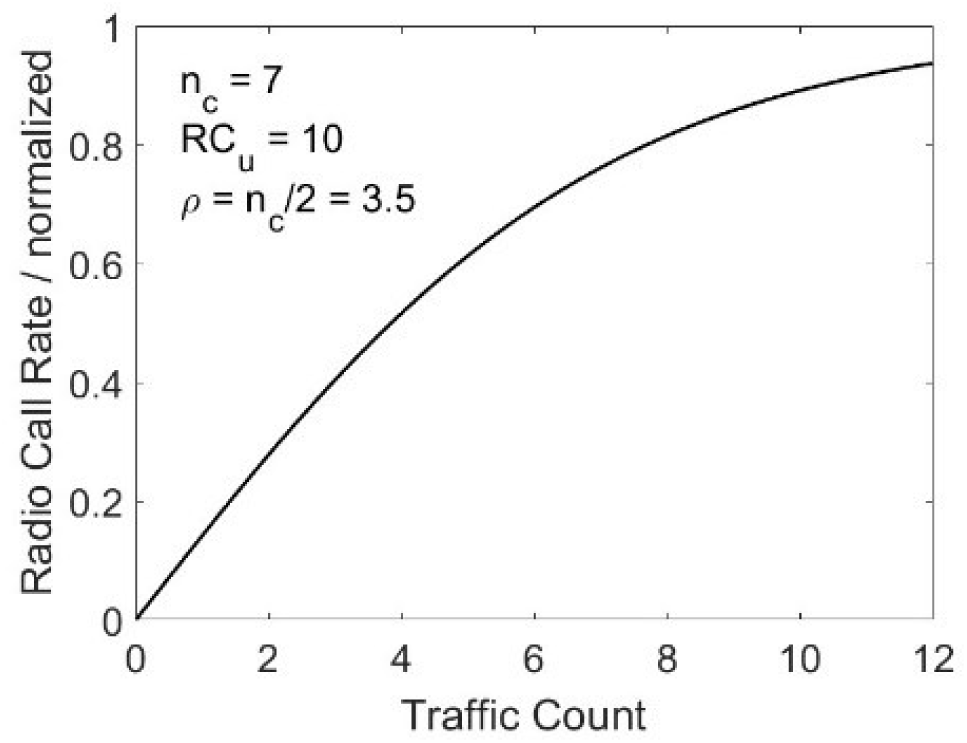
Theoretically predicted rate of normalized radio calls R(n)/R_u_ vs. traffic load. For the asymptotic value RC_u_ (or R_u_) we use as prior estimate of 10 calls/2 min. Scaling parameter estimate ρ calculated from linear extrapolation of slope at R(n = 0) := 0, using operational traffic capacity limit n_c_ ≈ 7 (≡N_Smax_ = sector traffic count; for details see text).

### 2.4. Cognitive Hill function as MWL(TL)-Model

In [1] and [2] we had shown that by combination of the logistic MWL- and TL-equations, (*4*) and (5) respectively, through replacement of the traffic load variable (n = ρ ln[(R - Δ_R_)/(R_u_ – R)] ) we can derive an ISA(RC)-power law with Δ_R_ := -R_u_, MWL-shift factor k = exp{μ/ν}= (I_u_-I_0_)/(I_0_-Δ). With exponent γ = ρ/ν the normalized ISA-MWL I(R)/I_u_ := p_I_( R) for bias Δ = 0 turns out as [1]:

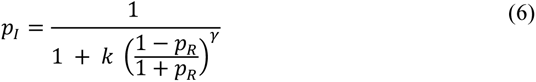

With 0 ≤ p_I_ ≤ 1. The transformation (R_u_ - R)/(R_u_ + R) = (1 - p_R_)/(1 + p_R_) := 1/S(p_R_) defines a new normalized cognitive stimulus (TL) variable S in the form of a contrast function, and 1 ≥ 1/S ≥ 0 for 0 ≤ p_R_ ≤ 1. It basically represents the ratio of remaining capacity for radio calls (R_u_ - R) to the sum of actually required (R) and maximum available (R_u_) cognitive TL-resources. In the following section 2.5 we show that equation (6) has the same form as the classical Hill function of hemoglobin-oxygen (HbO_2_) saturation in chemical equilibrium [3] [38]. For better comparison between cognitive and biochemical Hill function the logistic shift factor k can be replaced by S_0_^γ^ so that for S = S_0_ equ. (6) determines half maximum of the sigmoid p_r_ (S_0_) = 0.5. If we replace p_R_ by equation (5) the dependence of TL-stimulus function on traffic load is obtained as

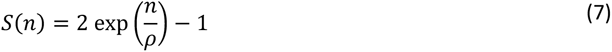

For Δ > 0 the cognitive Hill function has to take into account the (normalized) offset δ = Δ/I_u_ .

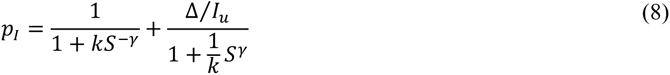

For Δ := I_d_ =1 the relationship k = S_0_^γ^ / (1 – 2δ) = 5/3 S_0_^γ^ holds. Because typically k can be very large (I_0_ – Δ) << 1) the power law MWL-bias approaches the bias Δ of the logistic I(n) characteristic. Figure 3 depicts theoretical predictions of the cognitive Hill model-equation (8) for Δ = 1, μ := n_c_ ≈ 7, ρ ≈ n_c_/2 = 3.5, and selected plausible intercept values {I_0_}:= Δ + {0.1, 0.25, 0.4}, with normalized cognitive TL and MWL variables (p_R_, p_I_).

**Figure 3:**
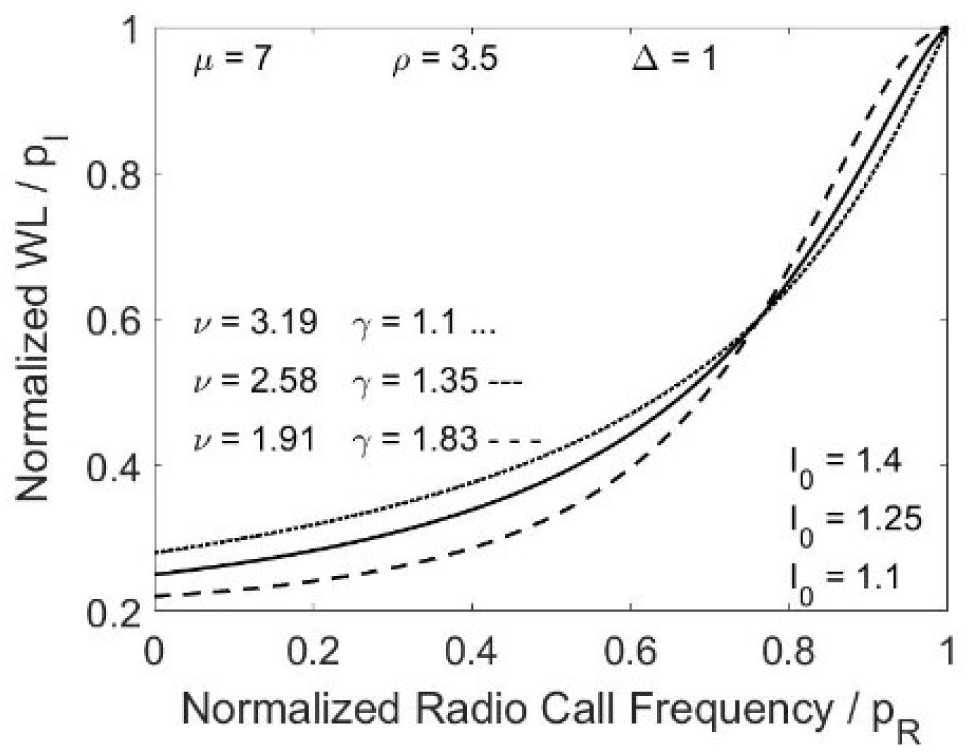
Power law curves of Hill-function (8) in normalized form p_I_(p_R_). Three intersection values (I_0_ = left endpoints of curves), corresponding to derived exponent values γ = ρ/ν, using {Δ=1, μ := n_c_ = 7, ρ ≈ n_c_/2 = 3.5} and logistic scaling ν (Δ, I_u_, μ, I_0_). For details see text.

The range of derived scaling parameter values ν = μ / ln(k) with k = (I_u_ – I_0_)/(I_0_ – Δ)) includes the theoretically estimated exponent γ = ρ/ν ≈ 1.5, the order of magnitude in agreement with Hill [3] and Stevens law exponent [19] [20] [15] (see below equation (9)). Extreme values of p_I_(p_R_) are for p_R_ = 0: p_I_(0) = Δ/I_u_ ≈ 0.2 (because k >> 1) and for p_R_ = 1 as expected: p_I_(p_Ru_) = 1. A close relationship between Hill-function (6) and Stevens’ stimulus – response power law was derived in [1] for ISA-MWL as dependent on RC-TL, through transformation of I(n) into P = (I - Δ)/(I_u_ - I). The new MWL variable P(I; Δ, I_u_) may be interpreted as ratio of required cognitive resources to remaining resources (I_u_ – I) that transforms (6) into the classical Stevens law [15] [16] [44]:

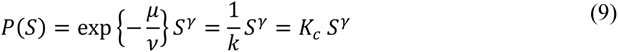

S(p_R_) as transformation of the logistic task load variable p_R_(n) serves as new independent cognitive TL variable for power law based MWL-data analysis and it is a mediator between the environmental load (simulation variable n) and subjective response. Because lim(S(p_R_ → 1)) = ∞, the graphical representation of p_I_ vs. S(p_r_) requires an upper p_R_-limit (maximum possible rate of radio calls as prior knowledge), e.g. S(R_max_ := 9 calls/2 min) = 19. In *Figure 4* equation (8) is shown in the standard inverse hyperbola form p_I_ vs. S(p_R_) of the Hill function, modified by the small bias term with Δ/I_u_ = 0.2 in equ. (8).

**Figure 4:**
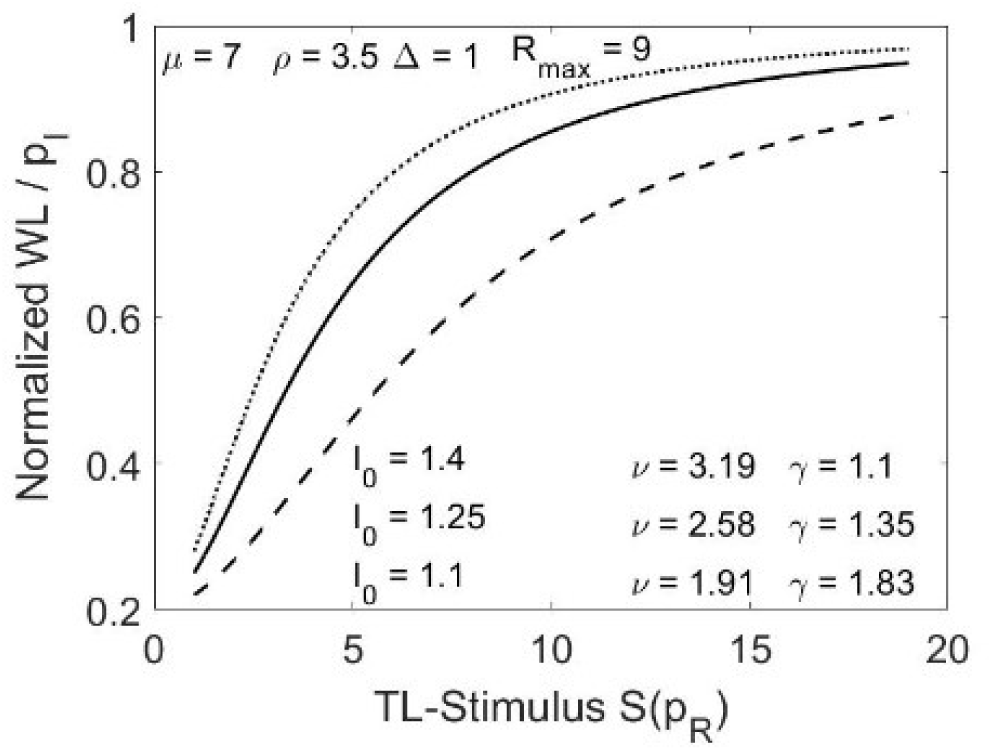
Hill function Equation (8) depicted in the standard inverse hyperbola form. Transformed abscissa stimulus scale S(p_R_). Same parameter values as Figure 3. For details see text.

The slope at half maximum, i.e. normalized MWL-increase with TL-stimulus variable (transformed radio calls rate S(R)) is derived as dp_I_/dS := p_I_’ = γ / (4 S_0_) ≈ 0.03 … 0.05, using γ = 1.5, S_0_ = k^1/γ^, k(I_u_, I_0_, Δ) = 19…39.

### 2.5. Equivalence of behavioral power law of cognitive load and Hill’s HbO2 saturation function

Based on research by Barcroft et al. from the Cambridge physiology laboratory (e.g. [45] [46]), Hill in a short 1910 publication formalized the fraction of Hb-receptor protein concentration that is bound by the O_2_- ligand [3] by means of the mass-action law (e.g. [47], see Discussion section 4) as:

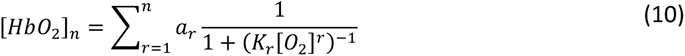

The weighted sum over inverse hyperbolas at the right side with exponents r = 1 ….n takes into account Hb-aggregates with n > 1. This means that the mass-action law for Hb_r_ + r O_2_← →Hb_r_O_2r_ equilibrium predicts a sum of saturation curves with integer exponents r = 1, 2, …, n with contribution a_r_. Hills fit to experimental data of [HbO_2_]_r_ saturation (solution prepared by Bohrs method, e.g. [48]) using n = 2 provided for r = 1, 2: K_1_ = 0.125, K_2_ = 0.0011 with weighting a_1_ = 0.38, a_2_ = 0.62. In fact it is textbook knowledge now (e.g. [49]) that the Hb-polypeptide is structured into n = 4 subunits, each one containing an Fe-binding site for O_2_-ligands, with the Bohr-effect (e.g. [48]) inducing self-organized, CO_2_-mediated enhancement of ligand binding through Hb- conformation change after the first O_2_ attached to a subunit (e.g. [50]).

Due to experimental difficulties in determining the parameters of equation (10), Hill proposed a simplified function for the equilibrium concentration R of the bound hemoglobin-oxygen complex [HbO_2_]_m_ as dependent on the O_2_ – ligand concentration L:

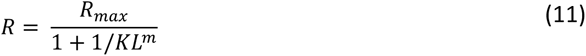

where R_max_ = total receptor concentration, and m > 1 assumed as a kind of average number of aggregated Hb molecules with Fe-attachment sites for O_2_ molecules. Ligand concentration L_0_ = L(R := R_max_/2) at half maximum of receptor-ligand complex is related the equilibrium dissociation constant with K = 1/L ^m^ [3] [38]. It quantifies the thermodynamics of chemical binding (e.g. [47]) between Hb and O_2_ via the Free Enthalpy difference ΔG^0^ = -RT ln(K) for the Hb + O_2_ ←→ HbO_2_ ligand association-dissociation equilibrium reaction under standard conditions [45]. Besides the above results obtained with equation (10), Hill showed in [3] a remarkably good fit to experimental results with the simplified “Hill function” (11): K = 0.0146 and exponent m = 1.4, where the deviation from 1 is due to simultaneous O_2_-binding to more than one of the four Hb-binding sites.

Evidently the Hill function is equivalent to our power law of cognitive load (equation (6) and (8) with separate bias term). This is not surprising due to the fact that both describe effects of resource limitation: the differential equations for time dependent changes of receptor and ligand concentrations as derived from the mass-action law ( [45]; e.g. [47] for basics) are closely related to the logistic functions of MWL and TL variations with environmental load n. It is easy to see that if we set for Δ = 0 in Equ. (6) k := k_0_ = S ^γ^ (first term in equation (8)) we get p_I_ (S := S_0_) = 0.5 so that S_0_ = k_0_ ^1/γ^ = exp(μ/ν) ^1/γ^ corresponds to the parameter L_0_. The ISA(n) shift-constant k = exp{μ/ν} = (I_u_ – I_0_)/(I_0_ – Δ) in equ. (*4*) and (6), (8) corresponds to the inverse of the dissociation equilibrium constant K (i.e. k = 1/K = 64.5). As theoretical prediction for the two parameters of the cognitive Hill-function we obtain with biased equation (8) (Δ = 1) the exponent estimate γ = ρ/ν = 1.1 - 1.8, for plausible assumptions for LRL-model parameters (see Figure 3 and Figure 4): I_0_ = {1.1, 1.25, 1.4}, {μ := 7 (see section 2.2), ρ = 3.5 (see section 2.3), ν( I_0_, I_u_, Δ)}. These assumptions yield cognitive dissociation constant K_c_ = 1/k = exp(-μ/ν) = {0.025, 0.068, 0.11}, anyway << 1. Also the normalized slope at half maximum of HbO_2_- saturation curve equ. (10) (R’/R_max_ = m /(4 L_0_) = m K^1/m^ / 4) = 0.017) compares reasonably well with our theoretical prediction for the cognitive power law derivative at half maximum p_It_’ = 0.03…0.05, again the same order of magnitude as the biochemical value (see section 2.4). These rough theoretical estimates of cognitive Hill parameters based on prior knowledge and plausible assumptions for the LRL-parameters turned out to be surprisingly close to the biochemical parameters of Hills’ classical function. Consequently, the theoretical predictions suggest a comparison with a corresponding analysis of recent experiments [1] [2] [4] [37] which provided initial evidence for the cognitive resource limitation hypothesis and the LRL-model of subjective cognitive load.

## 3. LRL-model based experimental Hill parameter estimates

In what follows we compare the regression based estimates for the behavioral Hill function parameters γ = ρ/ν and k = exp(μ/ν) = (I_u_ -I_0_)/(I_0_ – Δ) with Hill’s classical parameters of hemoglobin-oxygen saturation (saturation exponent m (vs. γ) [3] and HbO_2_ equilibrium dissociation constant K (vs. 1/k)), based on independent data sets from three ATC-HITL simulation experiments published in [1] [2] [32] and [4] [37]. With focus on the Hill function in the present work, the data sets were analyzed in the sequence of increasing agreement between cognitive and biochemical parameters and not in the sequence of their publication date because the original goal of the HITL-simulations with experienced ATC-domain experts (air traffic controllers, ATCOs) as participants was quantification of mental workload for various ATC-scenarios with different subjective and objective measures using conventional (linear) statistical analysis. The biased resource limitation (LRL) model for advanced data analysis with usage of prior information on parameter values was developed years after the experiments.

While experiments 1 [1] [32] and 2 [4] [37] had been realized through ATC-HITL approach-sector traffic simulation, experiment 3 had generated the first MWL and TL data [2] for the new Remote Tower Center work environment [51]. To my best knowledge the LRL model was the first one that theoretically predicted logistic dependencies of subjective MWL and objective TL on traffic load, and by means of variable transformation provided a MWL(TL) Hill function with direct relationship to Stevens’ power law. The present extension of our initial LRL-model in [1] [2] provides a theoretical foundation also for the results of Lee et al. in [4] [37] who to our best knowledge were the first to propose a heuristic four-parametric sigmoid MWL-model and determined parameter estimates through nonlinear regression analysis.

section 3.1 reviews briefly experiment 1 that had provided initial evidence for the cognitive resource limitation hypothesis [1] [32]. MWL- and RC-model parameters had been estimated through logistic regression with equations (*4*) (5) without ISA-MWL bias (equ. (*4*): Δ = 0). The approach traffic simulation data set of Lee et al. in experiment 2 ( [4] [37], section 3.2) with the 7-level ATWIT MWL-scale [12], are comparable also to our second data set (experiment 3, [52] [2], section 3.3). Like experiment 2 also experiment 3 provided a larger number of independent traffic load data points as compared to experiment 1 (with only four well defined traffic load values), so that in addition to the zero bias analysis (Δ = 0) we applied in section 3.3 also the extended LRL model regressions with Δ = 1 which improved the Δ = 0 estimates and confirmed the heuristic sigmoid four-parameter estimates [4] of Lee et al. in section 3.2.

For the LRL-model parameter estimates by means of nonlinear regression analysis we used Matlab®- nlinfit with robust option (bi-square residuals weighting) for reducing outlier effects. Details of the three experiments are briefly reviewed in the Methods overview of the supplement that also contains tables with the preprocessed data for experiments 2 and 3. For the latter extended data sets (different work conditions) are are available in the book chapter [2]. Experiment 1 data sets may be downloaded from [1].

### 3.1. Hill function parameter estimates from approach traffic HITL-simulation [1]

The ATC- HITL simulation of our first data set (experiment 1, [1] [32]) was mainly aimed at validating a new objective neurophysiological method of EEG based MWL measurement through quantifying the discriminability of ATCOs MWL-response. Four traffic flow values served as independent (environmental) variable, with increasing number n of aircraft (AC) per time interval in the sector with 25, 35, 45, 55 AC/hour, and n = n_c_ = 45 approximating minimum DC-separation (capacity limit) with typically 3 nm distance. They generated low, medium (typical for operational situations), high, and excessive task load, determined by radio communication between (pseudo) pilots and ATCO at the standard approach-sector radar-display workplace. Due to the low number (= 4) of independent variable values a simplified model with pre-selected ISA MWL-bias Δ := 0 = lower asymptote of the logistic sigmoid equ. (*4*) was used for minimizing number of logistic and power law regression parameters to one (μ for ISA, ρ for radio calls frequency (equation (5)), and γ = ρ/ν for Hill function (equations (6), (8)) and Stevens power law with ν = μ / ln(4) respectively). As a plausible value for this analysis we defined intercept ISA(zero traffic) = I_0_ := 1 = I_d_ (lowest MWL ISA-scale value ), with sigmoid shift factor k(Δ=0) = (I_u_ -I_0_)/I_0_ = 4 (see section 2.2). These assumptions determine the theoretical estimate for cognitive (behavioral) Hill constant K_c_ = 1/k = 0.25, an order of magnitude larger than Hill’s classical equilibrium dissociation value K = 0.014 [3]. In general however, we expect Δ =1 as lower asymptote, due to lower ISA and ATWIT scale limits I_d_ = 1 (see experiments 2, 3, following sections). Because in general intersection I_0_ > I_d_ and I_0_ – Δ << 1 with I_u_ - I_0_ ≈ 4, the shift factor k = 4 represents a lower limit and usually k >> 1 so that the behavioral estimate for K_c_ = 1/k << 1/4 is predicted, in agreement with Hill’s original K-value. LRL-model based parameter estimates with Δ = 0 described nearly linear MWL(n|μ) increase over the simulated range of traffic flow n [32] as predicted in the dashed curve of Figure 1.

A corresponding regression analysis for the radio calls task load data RC-TL(n; ρ) (averages across repeated measures and participants under standard task conditions) using equation (5) was published in [1]: ρ = 20.9 (±SE=0.3) calls/h. The power law regression with equation (6) of ISA-MWL(RC-TL) data provided the (cognitive) Hill-exponent estimate γ = 0.79 (0.01), with T-test p(T) = 6.4 10^-4^ (T = 70), about half as large as Hill’s biochemical value m = 1.4 obtained with his HbO_2_ saturation equation (11). The same (lower) value was obtained from our data via derived relationship γ = ρ/ν in equation (6) using the logistic regressions for ρ and ν = 26.3 AC/h with model equations (*4*) and (5). Anyway this γ-estimate is of the same order of magnitude as the biochemical estimate m. Analysis of the following two experiments shows the improved agreement by using the more realistic MWL-bias Δ := 1 as lower asymptote (extended LRL-model).

### 3.2. Hill function parameter estimates from approach traffic data of Lee et al. [4] [37]

For these MWL- and TL data of Lee et al. [4] [37] (extracted from Figure 4 in [4], see Supplement) obtained with the 7-level Air Traffic Workload Input Technique (ATWIT, [12]), the focus was on the transition between manageable and unmanageable traffic load. In contrast to experiment 1 with four well defined traffic flow values n_1_ … n_4_ (AC/h) as independent variable [1], the HITL-simulation of Lee et al. with four experienced ATCOs was designed for a traffic load continuum with Controller-Pilot Data-Link Communication (CPDLC, short “DL”) and Decision Support Tools (DST) to reduce workload, compared to radio communication. As a rough classification used for TL-analysis (see below), TL-ranges were characterized as “manageable”, “high”, and “unmanageable”. With regard to model based fitting of the subjective MWL-data Lee et al. compared heuristic linear, exponential and sigmoid relationships between the empirically observed cognitive MWL-ratings and managed traffic load. In this study of HITL ATC-control in approach sectors of three different US-airports a heuristic 4-parameter sigmoid-curve provided the best fit results to the ATWIT-MWL data:

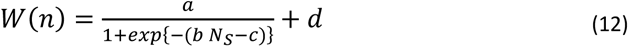

with N_S_ = sector traffic count as independent variable. In what follows we use W(N_S_) instead of I(n) for the subjective MWL-characteristic only with relationship to the heuristic model of Lee et al.. Correspondence to Verhulst equ. (3) parameters is given by: a := α/β, b := α, c := -(αs + ln(β)), d := Δ). The heuristic sigmoid parameters (a, b, c, d) of W(N_S_) can be interpreted in terms of the LRL-model function I(n)-parameters (equ. (*4*)), with (cognitive) load sensitivity (scaling) parameter ν, shift (translation) parameter μ, lower and upper asymptotes respectively (Δ, I_u_ (≡ W_u_) via: a + d = I_u_, b = 1/ν, c/b = μ. The authors concluded that within the continuous W(N_S_)-scale the subjective workload is categorial, where sudden jumps from category “manageable” to “unmanageable” workload can occur. The authors also emphasized, that from an operational point of view the transition from manageable (ATWIT < 5) to unmanageable workload (ATWIT ≥ 5) is of importance, at critical traffic load N_Smax_ (= “monitoring alert parameter” MAP). It appeared somewhat unexpected that the fitted upper sigmoid asymptote estimate was obtained by Lee as <W(n)>_u_ = a + d ≈ 4.7 for N >> N_Smax_ = MAP =18, instead of lim W(N_Smax_ → ∞) = W_u_ = 7. However, only a few subjective MWL reports > W(n) = 5 were monitored for N_S_ > MAP. Maximum sector traffic count N_Smax_ = D/d_min_ with d_min_ ≈ 3 – 5 nautical miles (nm) depending on AC-size, so that typically D ≈ 54 – 90 nm (see section 2.1). Using typical speed v = 120 -150 (nm/h) during approach, N_max_ may be translated into average traffic flow estimate n = N_S_ v/D = v/d_min_ = 40 – 50 (AC/h), in agreement with the “high” traffic flow limit (n_3_ = 45) of experiment 1. Lee et al argued that during overload situations (N_S_ > N_Smax_, i.e. communication task load DL-TL(N_S_) approaching upper asymptote) ATCO’s strategies change in order to avoid unmanageable traffic [4] (see also [30]). This was assumed to explain the observation of effective asymptote W_u_ ≈ 5 for traffic exceeding N_Smax_ ≡ MAP ≈ 18 AC, so avoiding the theoretically expected unmanageable upper ATWIT range, in favor of “safe” traffic control conditions.

Figure 5 compares the heuristic 4-parameter sigmoid fit (a, b, c, d: dashed curve) to the repeated measures data averages for each traffic load value (maximum traffic count during 5 min ATWIT-recording time intervals) obtained with the heuristic Lee et al. model, with the 3-parameter fit estimates (W_u_, ν, μ; solid line) using our LRL model equ. (*4*). Instead of Δ := 0 in the first experiment (section 3.1: zero ISA-MWL bias), we use here Δ := W_d_ = 1 by plausibility as prior information, i.e. lowest ATWIT reporting for vanishing traffic load. The figures depict averages across repeated simulation data for a single airport (a), and averaged across the three airports (b).

**Figure 5:**
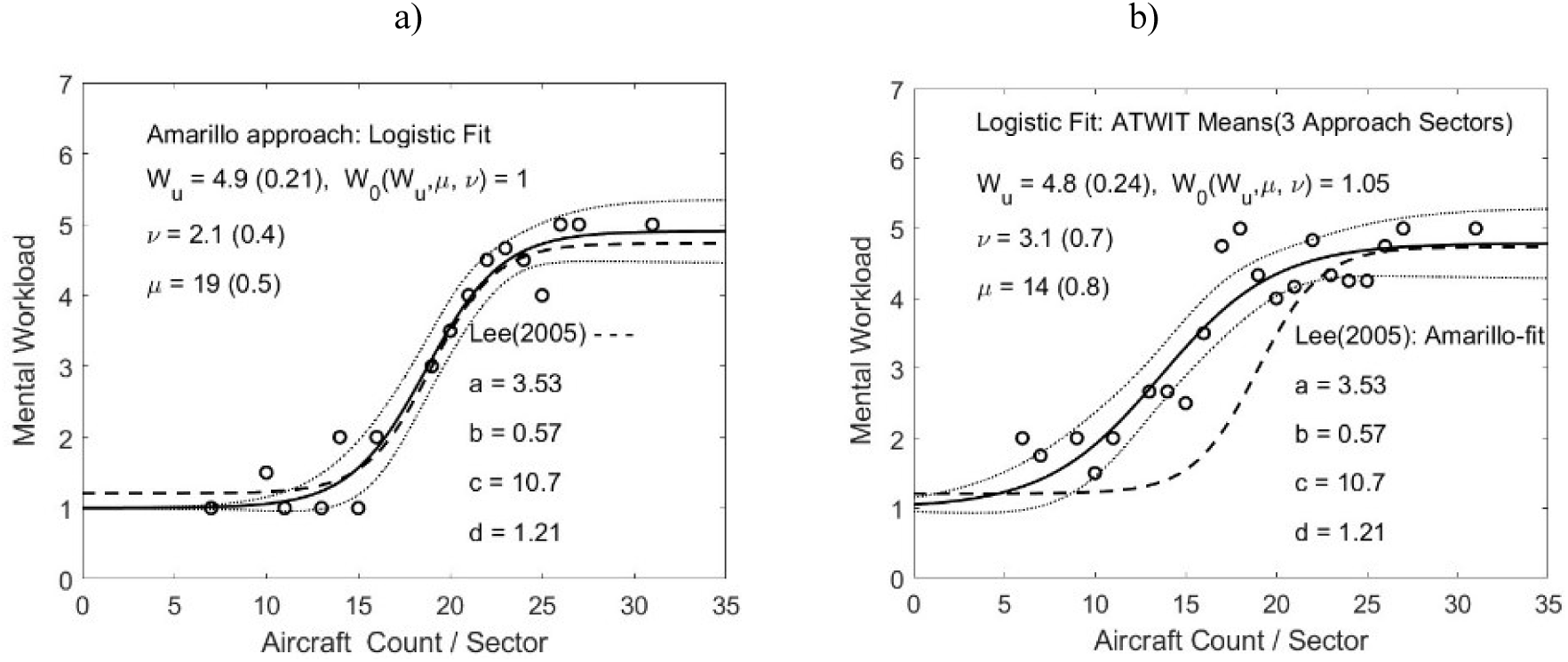
Approach sector control simulation with ATWIT-WAK MWL-data (5 min reporting intervals). Circles: averages of repeated WAK-reports, together with logistic regression with LRL model (solid curve). a): Amarillo approach. Dashed sigmoid: parameters from fit with equ.(12), (a, b, c, d) = (3.53, 0.57, 10.7, 1.21; specific sector monitor alert parameter MAP ≈ c/b = 18.8 ≈ μ = 19(0.5). Solid curve = logistic 3-parameter fit using equ.(4) with dotted curves for 95%-CI; parameter estimates with (SE): shift μ = inversion point, scaling ν = sensitivity parameter, W_u_ = estimate of upper asymptote; Δ := 1 as prior information = scale limit W_d_ , intersection calculated via W_0_(W_u_, ν, μ) > W_d_. b): ATWIT averages of MWL-data across Ardmore, Wichita Falls, and Amarillo results (circles) with logistic regression (solid) and 95% CI; numerical parameter estimates like in a).

In agreement with the prediction (solid curve in Figure 1), the exponential growth of MWL W(N_S_) in Figure 5a) b) starts with a weak nearly linear increase of the slope W’ = dW/dN_S_ for low MWL (N_S_ < 15 < MAP = 18) and continues with a nearly linear WL-range for W(N_S_) around the inversion point (N_S_ = μ) at half maximum between asymptotes (slope maximum = (W_u_ – Δ) / 4ν) before W’ decreases to zero, i.e. W(N_S_) converges to (effective) W_u_ ≈ 5.

For the original analysis in [4] the authors reported a Pearson correlation of R^2^ = 0.84 (dashed curve; remark: we corrected one of the reported parameters with respect to the sign: c = +10.7; mistakenly -10.7 in [4]). Parameter estimates for both fits of the Amarillo approach data (Figure 5a), dashed and solid curves) exhibit agreement (within the given uncertainties (stderr SE)) if we replace the estimate d = 1.21 by Δ = W_d_ := 1 for our 3-parameter regression. Correspondences of our logistic parameter estimates (±SE) and those of [4] from 4- parameter fits are: W_u_ = 5 (0.2) ≈ a + d = 4.74 ≈, ν = 2.1 (0.4) ≈1/b = 1.8, μ = 19 (0.5) ≈ c/b = 18.8. With the three parameter estimates the intersection W_0_ ≡ W(n=0) (≡ I_0_ in equ. (*4*)) can be calculated via W_0_ = (W_u_ + kΔ)/(1 + k) = (5 + k)/(1 + k) ≈ 1, using k = exp(μ/ν) = 1.34 10^4^). The normalized slope of the sigmoid (i.e. maximum WL sensitivity) at the inversion point μ = 19 (AC count) from the Amarillo fit is dW_u_/dn/(W_u_ – Δ) =1/4ν = 0.12 (per AC). Comparing a) and b), the decrease of slope W’ ∼ 1/ν marks a less rapid transition to unmanageable load under increasing traffic N_S_ for b). For the average across the three sectors the slope decreased to 0.08 (per AC) with ν = 3.1 (±0.7), and with μ = 14, μ/ν = 4.52 and k = exp(μ/ν) = 91.8.

We use this latter (averaged) value for comparison with Hill’s dissociation constant K because despite the comparable parameter uncertainties in a), b) the larger intercept W_0_ for b) results in a more reliable k-value through k = (W_u_ – W_0_)/(W_0_ -1) due to a significantly larger distance (W_0_ – Δ) for b) than for the single airport result a). For the cognitive Hill function constant this LRL-model estimate yields K_c_ = 1/k = 0.011 that compares well with Hill’s dissociation equilibrium estimate K = 0.014 reported in [3].. Consequently the agreement of MWL parameter estimates (μ, ν, within CI) with the heuristic model based analysis of the Lee et al. supports the theoretical basis of the LRL model as well as the prediction of the Hill function equilibrium constant..

Estimate of Hill function exponent γ requires direct regression of corresponding ATWIT-MWL vs. TL data with power law equation (8), or alternatively of DL-TL vs. N_S_ data for estimate of ρ via tanh-function (5) and γ = ρ/ν according to theory. However, in contrast to MWL(N_S_) in [4] only three communication-TL measurements regarding number of data link (DL) contacts (handoffs and clearances for ATCO 1 and 2) are reported for the three airport simulations in [37] (averages <DL> per 10 min intervals for corresponding average traffic counts <N_S_>), roughly discriminating between manageable (m), high (h), and unmanageable (u) traffic load. No data were published for low traffic. We use the most reliable data (according to Lee et al.) from the Ardmore approach simulation [37] (from Amarillo only h and u available, from Wichita Falls m and h are inconsistent): {(<N_S_>, <DL>)_m, h, u_} = {16.1, 58), (18.5, 72), (21.3, 80)}. The decrease of slope Δ<DL>/Δ<N_S_> between the two steps from m to h := 14/2.4, and from h to u := 8/2.8 indicate the theoretically predicted resource limitation effect according to equation (5) and Figure 2: <DL> / D_u_ := y = tanh(N_S_/2ρ). Because the maximum DL(N_S_) = 80 can be assumed close to the resource limit we set asymptote DL_u_ := 85 handover-calls (per 10 min) which is somewhat larger than the theoretical prediction of equation (5) (400 calls per hour) depicted in Figure 2, section 2.3 (not surprising due to CPDLC-automation in the Lee et al. simulation experiment).

An estimate for the TL-parameter ρ is then obtained with the inverse of the theoretical saturation sigmoid (equ.(5)), i.e. the Atanh-function x= N_S_/2ρ = ln{( 1 + y)/(1 - y)}. After normalization the three data points from [37] {x_m,h,u_ = <N_S_> /2ρ , y_m,h,u_ = D_u_ <DL>} yield scaling parameters via ρ = <N_S_ >/ 2x: ρ_m,h,u_ = {4.84, 3.69, 3.06}with mean <ρ> = 3.9 (20% uncertainty ±0.8 corresponding to lower/upper mean). With above sensitivity parameter estimate ν =3.1 (3 AP-average ±0.7, Figure 5) the cognitive Hill exponent <γ> = <ρ>/ ν = 1.3 (±0.3), in surprising agreement with the biochemical Hill exponent m = 1.4 in [3].

### 3.3. Hill function parameter estimates derived from RTC- HITL simulation [2]

The third HITL-data set with quasi real-time subjective ISA-MWL recording (experiment 3 with 12 experienced domain experts; see supplement for methods-info and preprocessed data table) was obtained with Remote Tower Center (RTC) simulations in 2011 [2] ( [51]). It addressed the usability-quantification of the new RTC work environment for tower controllers through statistical standard methods ( [52], ANOVA and linear correlation analysis). Again it should be emphasized that also this experiment like experiment 1 (section 3.1) originally was not designed for advanced (nonlinear, LRL-) model based MWL-data analysis. As before we compare LRL based behavioral power law parameter estimates {k, γ} of the data obtained under standard work conditions, with biochemical Hill parameters {K, m}. Like in the first experiment ( [1], section 3.1), in [2] an initial set of logistic parameter estimates {μ, ν} and {ρ, R_u_} for the ISA(n)-MWL and RC(n)-TL functions (*4*)(5)respectively was obtained without ISA-bias, i.e. Δ := 0, I_0_ := I_d_ = 1 as prior assumption, with upper ISA limit I_u_ := 5 and TL-origin RC(n=0) := 0. In addition, for a direct comparison with the MWL-results of Lee [4] (section 3.2: experiment 2) we performed in the present work a new LRL-model regression analysis with ISA- bias Δ = 1 (extended model) instead of Δ = 0 (no bias). This appeared appropriate because in contrast to the first experiment [1] with repeated measures at four well defined values only of the independent traffic load variable n (AC/Δt; low, medium, high, excessive), we had in [2] a kind of sector-traffic count continuum in the range 0 < N_S_ < 10, comparable to [4]. Our preprocessed raw data (repeated measures averaged for each N_S_ and across participants) together with logistic regressions for parameter estimates of ISA(N_S_) and RC(N_S_) are depicted in Figure 6.

**Figure 6:**
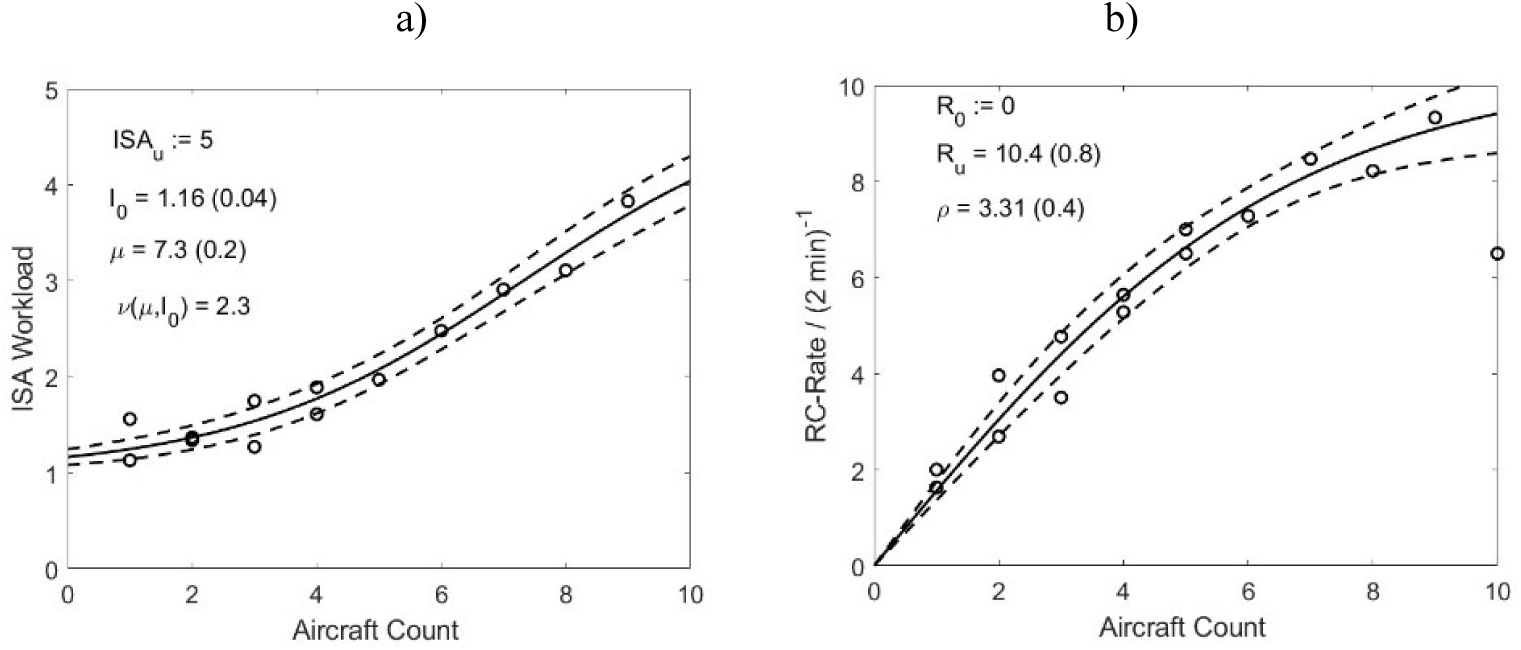
Results of RTC-HITL simulation experiment. a) Subjective ISA(N_S_)-MWL with 2 min MWL-reporting interval, and b) objective RC(N_S_) radio calls per 2 min (averaged across participants and data cases (2 min intervals)) as dependent on environmental traffic count N_S_. Robust logistic 2-parameter lsq. fits (solid lines) with 95% CI (dashed). ISA estimates I_0_, μ, using prior information for upper and lower asymptotes, I_u_ and Δ = I_d_ =1 respectively; ν(I_0_, μ) derived from estimates. RC estimates R_u_, ρ, with offset Δ_R_ := - R_u_. i.e. R(N_S_=0) := 0. Bi- square residuals weighting for outlier suppression. Data cleaning: outlier removed at ISA(N_S_ = 5), i.e. high uncertainty due to < 5 measurements /cases per N_S_-reporting / recording interval used for averaging.

Clearly both ISA-MWL and RC-TL characteristics exhibit pronounced nonlinear dependence on traffic count N_S_ and are fitted well by the LRL-model. Data show a range of very low, and similar to experiment 2 [4], a few excessively high MWL and RC-load values near the asymptotic scale limits. This is also depicted in Figure 7 that illustrates the direct power law regression analysis of the ISA(RC) data with the cognitive Hill function. As expected the theoretically predicted cognitive load characteristics ISA(R) of Figure 3 (equation (8) with S(R/R_u_) and Hill’s inverse hyperbolic saturation curve [3] ISA(S) of Figure 4 (equation (8) with RC-TL axis transformed into normalized scale S = (R_u_ + R)/(R_u_-R)).

**Figure 7:**
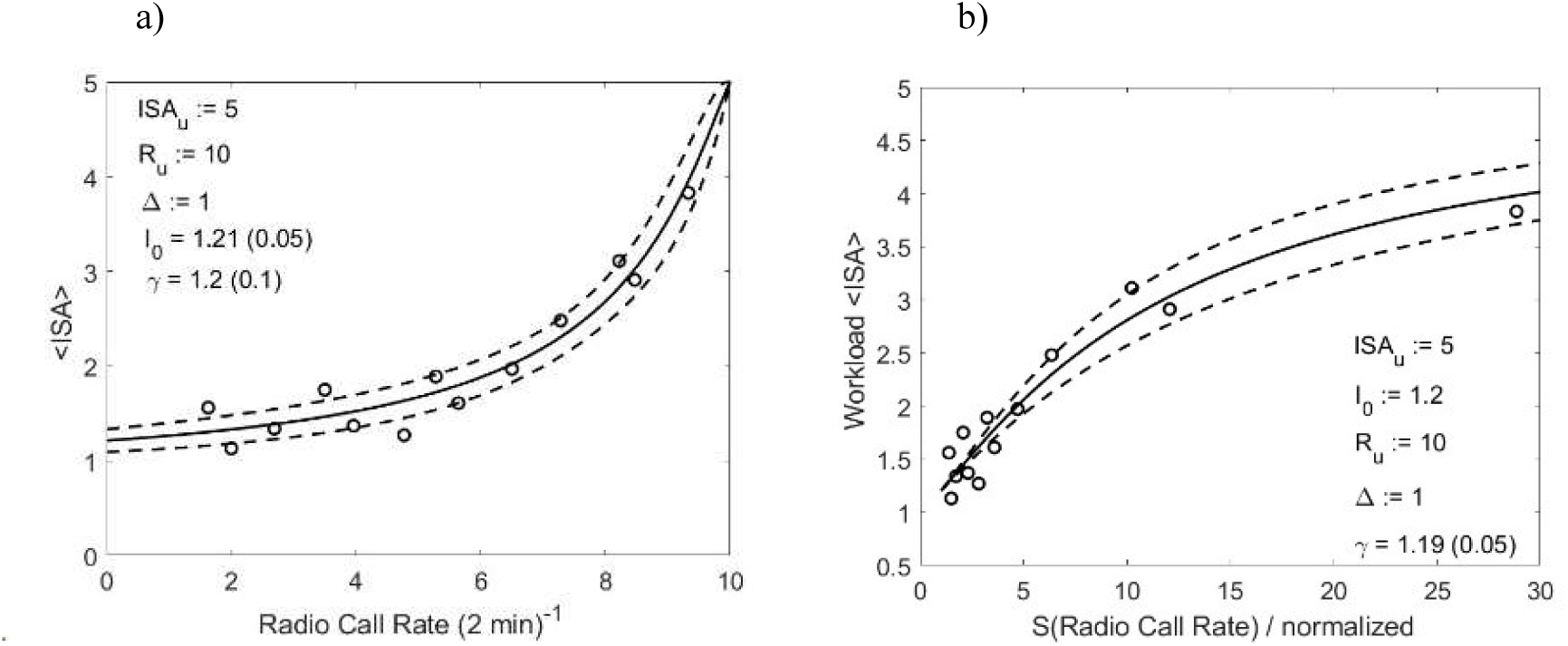
a) ISA-MWL data (averaged across 12 participants) as dependent on RC task load variable (radio calls per 2 min), with robust 2-parameter ISA(RC-TL) lsq. fits (equ. (8) with S(R/R_u_), solid line with 95% CI (dashed); inset: parameter estimates (±SE) for intersection I_0_, exponent γ including prior knowledge for I_u_, R_u_, Δ. b) The same data fitted with equ.(8) depicting asymptotic cognitive resource limit by using transformed TL-scale for ISA(S). Solid line: 1-parameter fit with 95% CI (I_0_ := 1.2 assumed as additional prior information).

For cognitive parameter estimates with the given data sample the Hill function with linear RC scale appears more appropriate as compared to the transformed S(R)-scale, the latter showing concentration of data in the low S-range. As expected, within uncertainties the cognitive Hill function fit shows the same exponent γ estimate as the value calculated from the logistic regressions via γ = ρ/ν. The same is true for shift factor k = (I_u_ – I_0_)/(I_0_ – Δ) = 19. Results of both sets of LRL-parameter estimates (with Δ = 0 and 1) are shown in Table 1.

**Table 1:** LRL-model based logistic regression analysis of MWL and TL data from RTC-HitL-simulation. Standard work conditions (1 ATCO per airport). 2-parameter lsq.- estimate of ISA(N_S_; I_0_, μ) with calculation of sensitivity parameter ν(I_0_, m). One-parameter estimate of RC(n; r) with prior assumption R_u_ := 10. Derived cognitive Hill-function parameters k, (1/k = K_c_), γ, for comparison with biochemical Hill-parameter values [3]: exponent m = 1.405, equilibrium constant K = 0.01405.

| Logistic 2- parameter regression (±SE) | | | | | 1 param. regression | | Hill function parameters | |
| --- | --- | --- | --- | --- | --- | --- | --- | --- |
| $\Delta$ | $I_0$ | $\mu$ | $v(I_0, \mu)$ | $k(\mu, v)$ | $R_u$ | $\rho$ | $\gamma$ | $K_c = 1/k$ |
| 0<br>Ref.<br>[2] | 0.83 (0.1) | 6.1 (0.4) | 3.6 (0.5) | 5.4 | 10 | 3.1 (0.1) | 0.70 (0.1) | 0.19 |
| 1 | 1.16 (0.04) | 7.3 (0.2) | 2.3 (0.2) | 23.9 | 10 | 3.1 (0.1) | 1.35 (0.1) | 0.042 |

For RC(n ; ρ) we assumed R_u_ = 10 as prior knowledge that improved the ρ-uncertainty significantly. It turned out that parameter estimates with Δ = 1 exhibit lower (rel.) uncertainties as compared to Δ = 0 [2] due to improved LRL-modeling of the extreme low and high traffic load ranges (asymptotic, exponential character). Like for the Lee et al. parameter estimates with Δ ≈ 1 in the previous section, the estimate of the cognitive Hill constant K_c_ = 1/k(Δ=1) again is of the same order of magnitude as the equilibrium dissociation constant K = 0.015 in [3]. While this agreement is not quite as good as in experiment 2, the cognitive Hill exponent γ agrees perfectly (within SE) with Hills estimate of the biochemical exponent m = 1.4.

## 4. Discussion

In the present work we provide evidence for the equivalence of subjectively experienced mental work load (MWL) as dependent on task requirement, and hemoglobin-oxygen (HbO_2_) saturation which can both be formally described by a Hill (power law) function. While the classical Hill equation was derived from the mass-action law (e.g. [47] [53]) of the chemical HbO_2_ ←→ O_2_ equilibrium [3], the cognitive Hill function is obtained by combination of (biased) Logistic Resource Limitation (LRL) models of subjective MWL and objective task load (TL). The specific LRL-functions of ATC-domain expert’s (quasi) online measured subjective MWL(n) and objective TL(n) characteristics with air-traffic load n as independent simulation variable were obtained as solutions of the nonlinear (1^st^ order 2^nd^ degree) Verhulst differential equation as most simple approach to formalize the effect of cognitive resource limits (e.g. processing capacity and working memory limit, [5] [6] [11] [54]). General solution is the logistic function represented by two-parametric sigmoid curves with exponential scaling ν (ρ for TL(n)) and sigmoid shift μ, or shift factor k = exp(μ/ν) (:= 1 i.e. μ := 0 for TL(n)). Interestingly, the cognitive Hill (inverse hyperbola [3] [38]) function (6) (8) may be transformed into the classical (stimulus – response) Stevens power law (9) of psychophysics. The LRL-model allows to combine objective TL and subjective WL sensitivity parameters (ρ, ν) into a single psychophysical (Hill and Stevens) exponent γ = ρ/ν. The theoretically predicted and experimentally confirmed order of magnitude (≈ 1) is consistent with the classical Hill-function exponent of Hb-Receptor O_2_ -Ligand saturation. The inverse of the logistic MWL shift factor(cognitive Hill constant) K_c_ = 1/k corresponds to dissociation equilibrium constant (e.g. [47]) *K* = *exp*(–Δ*G*^0^/*RT*) defined through the thermodynamics of Hb + O_2_ ←→ HbO_2_ binding with free enthalpy of formation ΔG^0^, gas constant R = 8.314 J/mol/K, and absolute temperature T(K), with initial quantification in [45].

From a biophysical point of view the resource limits are determined by the energy supply through hemoglobin (Hb) oxygenation ([HbO_2_]_m_, m = 1…4, e.g. [55] [49] [50]). For his classical biochemical function of oxy- hemoglobin saturation, Hill simplified the exact solution of the corresponding physico-chemical (1^st^-order 2^nd^- degree) mass–action law (weighted sum of four hyperbolas with exponents 1 ≤ m ≤ 4) through a kind of average single exponent for O_2_-binding to the Hb_m_-protein complex. He proved it sufficient for a surprisingly good fit the early experimental HbO_2_ saturation curves (see section 2.5). Consequently, from the perspective of cognitive resource limitation dynamics (Verhulst equation) and bio-chemical mass-action law for HbO_2_ receptor-ligand saturation equilibrium, it is no surprise that limitation of neural energy supply determines cognitive resource limits. It predicts a parametric correspondence between cognitive (k(μ, ν), γ(ρ, ν)) and biochemical (K ←→1/k, m ←→ γ) power law parameters (equations (6) (8) and (11) respectively).

In the present work MWL and TL data from three previously published air traffic control Human-in-the-Loop (HITL)-simulation experiments were re-analyzed [4] [1] [2] with ATWIT-WAK [12] and ISA MWL- measures [34] [13] for quantifying the theoretically predicted correspondence between the cognitive and bio- chemical Hill function parameters. Highly experienced domain experts (air traffic controllers) as participants had allowed for relatively low inter-individual scattering of subjective cognitive load measurements. These kind of laboratory conditions under nevertheless realistic work environments and task settings had provided unique conditions for validating the theoretically predicted logistic workload and task load characteristics.

Table *2* collects estimates of cognitive Hill exponent γ and constant K_c_ = 1/k obtained from theoretical predictions (column 2) and from regression estimates using LRL-model based logistic functions from the three different experiments. without ISA-bias (experiment 3, Δ := 0, columns 3, 5), and with bias (Δ = 1, experiment 2and 3 (new analysis of data in [2] [52]), columns 4 and 6 respectively) .

**Table 2:** Theoretical and experimental estimates of power law exponent γ and cognitive Hill constant K_c_ = 1/k. Values for bias Δ := 0 (initial model [1] [2]) and Δ = 1 (extended LRL model) as prior information, to be compared with biochemical Hill parameters m = 1.5, K = 0.014. Uncertainties (±SE(γ, K_c_)) via error propagation with (δρ, δμ, δν)). Columns 3, 5: Δ := 0 [1] [2], Columns 4, 6: Δ := 1. *For experiment 2 [4] [37] a systematic error δγ ≈ 0.25 > SE(γ) may be assumed due to systematic ρ-increase with N_S_.

| Cognitive Hill Parameter | LRL-Theory<br>$\rho := 3.5$<br>$\mu := 7$<br>$I_0 := 1.1-1.4$<br>$\Delta := 1$ | Experiment 1:<br>Approach-HITL data [1] [32] with LRL model regressions<br>$\Delta := 0$ | Experiment 2:<br>Approach-HITL data Lee et al. [4] [37] with LRL model regressions for $\Delta := 1$ | Experiment 3:<br>RTC-HITL data with LRL model regressions for<br>$\Delta := 0$ [2] | Experiment 3:<br>RTC-HITL data [2] with LRL model regressions for<br>$\Delta := 1$ |
| --- | --- | --- | --- | --- | --- |
| $\gamma = \rho/\nu$<br>($\pm SE$) | 1.8...1.1 | 0.70 (0.05) | 1.25 (0.14)* | 0.79 (0.01) | 1.35 (0.1) |
| $K_c = 1/k = \exp(\mu/\nu)$ | 0.026 ... 0.05 | 0.25 | 0.011 (0.012) | 0.25 | 0.043 (0.012) |

Table *2* shows that for the offset Δ = 1, the theoretical estimate of the Hill-/ Stevens cognitive power law exponent γ = ρ/ν (column 2) based exclusively on prior information, as well as estimates based on logistic regressions for ν and ρ (columns 4, 6) are consistently > 1. Within uncertainties they are in surprising agreement with Hills original estimate m = 1.4 in [3].

While the estimates of cognitive Hill exponents for MWL-bias Δ = 0 are consistently significantly lower than m for both experiment 1 and 3 (columns 3, 5), the associated cognitive (equilibrium) constant K_c_ = 1/k are fixed by prior model conditions as k = (I_u_ -I_0_)/(I_0_ – Δ) = 4 and indicate an upper limit, an order of magnitude larger than K. The data analysis with initial model bias Δ = 0 (i.e.no MWL bias) was selected in [1] due to the low number (= 4) of different external load (independent) variable values (n (AC/h) = 25 … 55 corresponding to (approach) sector count N_S_ ≈ n D / v ≈ 10 … 22, with v ≈ 120 knots, D ≈ 50). In contrast, the extended model with Δ = 1 was required for an optimized regression with a higher number of traffic load values, with Δ = 1 for experiment 2 [37] (K_c_ = 1/k = 1/92 = 0.011), and experiment 3 [2] (K_c_ = 1/24 = 0.04). Within uncertainty ranges the resulting cognitive K_c_-values again compare well with biochemical K = 0.014 in [3]. Under the hypothesis that subjective ISA-MWL reporting represents a true measure of cognitive resources and linked directly to HbO_2_-saturation (i.e. available neural energy), it appears plausible that Δ = 1 provides better agreement with the Hill coefficients than Δ = 0 because the HbO_2_-saturation scale starts at 0 % (ending at 100%) whereas the arbitrary ISA scale starts at I_d_ = 1. The latter value may be interpreted as representing basic mental load for ongoing low level task requirements, in addition to the variable traffic load simulation variable n.

It is straightforward to explain the increase of the cognitive Hill-exponent from γ < 1 for bias Δ := 0 in our previous initial analysis [2], to γ > 1 with the biased LRL-model using offset Δ = 1 as prior knowledge in [2], if we take into account that γ ∼ 1/ν ∼ slope I’ = dI/dn of the logistic ISA characteristic. We had shown in [32] that due to the only four discrete traffic load values of the independent variable the MWL-sigmoid was approximated reasonably well by a linear model. Under standard work conditions the MWL-characteristic with Δ := 0 (like in [1]) starts at intersection I_0_ := I_d_ = 1 with a slope (or WL-sensitivity) I’ = dI/dn = 4/(5ν) that is certainly larger than slope dI/dn (Δ = 1) ≈ 0 (like in [4]), due to the extended low-traffic range of the latter that shifts the sigmoid slope-inversion vale μ (transition coordinate in [4]) to the right (e.g. Figure 5). Consequently, for a given range of the environmental traffic load variable n (0 … n_max_ > n_c_) the latter I(n)-characteristic is forced into a steeper increase with maximum slope I’_max_(n = μ) = (I_u_ – Δ) / (4ν) if we assume for both cases the same cognitive resource limit I_u_ and a comparable critical environmental load value n_c_. So, if we require I’_max_(Δ = 0) < I’_max_(Δ = 1) ∼ 1/ν the WL-sensitivity parameter ν has to be significantly smaller for Δ = 1 as compared to Δ = 0, which together with the same TL-sensitivity parameter ρ in both cases, results in a significantly increased power law exponent γ = ρ/ν, in agreement with the observed parameter values in *Table 2* .

There are at least two reasons why the range of validity of postulated equivalence between cognitive and biochemical Hill functions should be limited. On the one hand the classical Hill function is an approximation to be used for simplifying data analysis, providing a kind of average across the four contributions to the saturation curve model equ. (10) of the four Hb-protein O_2_-binding sites. Also Hill in his classical article, besides the (K, m) parameter fit result for the reported dissociation curve used for the comparison with our LRL- model parameters (K_c_, γ) , provided a list of parameter fit pairs obtained from six different HbO_2_ saturation curves (from experiments with different salt solutions) that exhibited variations in the range (0.000257, 3.19)… (0.0453, 1.670)… (0.00427, 2.111).

On the other hand the HITL-simulations under roughly doubled task requirements (experiment 3: single ATCO controlling two airports) suggesting bias Δ > 1, revealed an increased tendency of work strategy switching of domain experts to avoid MWL overload under excessive TL resulting in deviations from the standard MWL(n), TL(n) curves (see discussion in [2] and Figures 9 and 10 therein). That is why it appears of interest to compare theoretical logistic and cognitive Hill-parameter predictions for increased task requirements (due to different work conditions) resulting in MWL-bias Δ > 1, with parameter estimates from corresponding experimental data as reported in [2]. In the latter experiment MWL-bias Δ = 2 was assumed for optimized LRL- based parameter estimates under conditions of roughly doubled (secondary) task requirements for low traffic load. Corresponding investigations are under way and will be reported in a follow-up paper.

## 5. Conclusion

The unexpected quantitative agreement between cognitive and biochemical Hill parameter values ({K_c_, γ} and {K, m} respectively) for MWL-bias Δ = 1 of the extended LRL-model justifies the hypothesis that within our given experimental conditions (realistic HITL-simulation with highly qualified and motivated ATC-domain experts as participants and quasi real-time MWL reporting) the subjective ISA and ATWIT MWL-reporting as dependent on the normalized task load in the form of a cognitive Hill-function (equ. (8) in section 2.4), could be interpreted as an (interoceptive) measure for the decrease of available energetic resources under increasing cognitive load due to HbO_2_ - saturation limiting the ATP production. This conclusion is based on the finding that the cognitive Hill function (initially derived in [1] as equation (7) for Δ = 0: equation (6) in the present work) is formally and quantitatively equivalent to Hill’s classical function in [3]. This equivalence may be taken as a mathematical support for results of embodied cognition research which have revealed the influence of energy consumption on decision making (i.e. Glycolyse, e.g. [40] [56]). Our theoretical approach also exhibits a relationship to Friston’s unified brain theory based on the free energy principle [24], and to the “selfish brain” theory of Peters e.al. [22]. In contrast to our non-linear logistic resource limitation approach, the latter is based on a linearized logistic supply-chain model [25] that formalizes the self-organized homeostasis of cerebral metabolism. and that is able to describe also stress related neural diseases under malfunction [57] . Consequently our hypothesis may be interpreted as follows: task load (TL) increase under increasing task-requirements involves increasing use of energy resources (i.e. Glycolyse) with corresponding increase of O_2_ transfer into the brain cells via HbO_2_ dissociation, enhanced by the blood CO_2_ concentration through Hb-protein conformation change (Bohr effect, e.g. [48]). Formalized by the Hill equation the ISA- and ATWIT-MWL(TL) report as subjective decision on experienced cognitive load level represents an interoceptive sensing of the remaining energetic resources through the direct correspondence between normalized task load function∼ (K_c_ S)^-γ^ and the function of blood oxygen (ligand) concentration ∼ (K L)^-m^. Moreover it is of interest that the close relationship between Hill and Stevens functions (see section 2.4) puts also the latter as classical (physical) stimulus – response power law on the fundamental biophysical/biochemical basis (section 2.5) of HbO_2_ saturation with Glycolyse and resource limitation by cerebral energy consumption. With regard to logistic MWL and TL LRL- model parameters (μ, ν) and ρ respectively, dissociation constant K via ΔG^0^/RT = - ln(K) relates the free enthalpy of HbO_2_ formation to the logistic MWL(n) sensitivity ∼ μ/ν = -ln(K_c_), and Hill or Stevens exponent m to the ratio of logistic MWL(n) vs. TL(n) sensitivities ρ/ν.

In order to improve the evidence for our hypothesis (or falsify it) additional data available in [1] [2] [52] and obtained under conditions with increased task requirements and corresponding additional MWL-bias are presently being analyzed with the extended LRL model. However, a specifically-designed new HITL-experiment would be required as an “experimentum crucis”, possibly including real-time measurement of physiological HbO_2_-saturation with advanced (fNIR) methods like in [10]. The range of the independent variables should be matched to the transformed stimulus variable-scale S(TL) of the cognitive power law and the participant sample should consist of experienced task-domain experts.

## Acknowledgement

I am indebted to Anne Papenfuss and Christoph Möhlenbrink who together with Michael Lange designed and organized experiment 3. A.P. provided the respective preprocessed raw-data used for testing the theoretical Hill- parameter predictions of the present work. She was co-editor of the 2^nd^ edition of the book “Virtual and Remote Control Tower” and co-author of a book chapter on model based RTC-WL-parameter estimates, and of an internal DLR-report on the initial data analysis of those early HITL-RTC-simulations.

## Supplement

### 1. Methods

Here we present a brief summary of the methods used in each of the previously published three HITL ATC- simulation experiments used for testing the (behavioral) cognitive LRL-model based predictions of the Hill-function parameters of HbO2-saturation as derived in the present paper. Details can be found in the referenced original publications. It should be mentioned that none of these experiments had been designed for testing the LRL model or the correspondence between cognitive and biochemical Hill function.

#### Experiment 1

The HITL approach traffic simulation with twenty-one well trained ATC-domain experts and controllers (ATCOs) took place in a realistic work environment with approach traffic radar-display [32] [1] and data link for simulated radio communication with pseudo pilots. It was based on four traffic scenarios for low (25), operational (35), high (45), and extreme (55) traffic flow n [aircraft per hour entering approach sector] as independent environmental load variable. MWL was recorded online in 2 min intervals using the discrete ISA- measure (I_d_ = 1 ≤ ISA ≤ I_u_ = 5) during the simulation runs of 20 min duration. Participants used a touchscreen for reporting their subjectively experienced mental load in quasi real-time. Simultaneous monitoring of radio communication between ATCOs and pseudo-pilots in a separate room provided the rate of radio calls [calls per hour] as objective (cognitive) task load (TL) variable. Each participant repeated eight scenarios with the different traffic load values in randomized order in two consecutive half days. Four of these scenarios included a non-nominal event (factor 2) after 10 min of simulation time to increase task load under the same traffic load for comparison of experienced MWL under increased stress. These were not evaluated for the present investigation that focused on standard work conditions with the single independent load variable n_i_, i = 1,…, 4. Logistic and Hill function based parameter estimates through nonlinear regression were performed for the averages across participants of MWL(ni) and TL(ni) data for each of the four standard traffic scenarios.

#### Experiment 2

The MWL and TL-data of HITL approach traffic simulation results of Lee et al. [4] [37] were re-analyzed with LRL-model based regression analysis. The measured discrete MWL-data were extracted from Figure 4 in [4]. In their experiment, each one of four professional air traffic controllers (ATCOs) with very high familiarity with their individual sector took part in the high-fidelity simulation (with advanced CPDLC data link between ATC and aircraft flight management system, replacing radio communication) of three US airports (Amarillo, Wichita Falls, and Ardmore). Traffic count NS in the sector as independent environmental load variable, and task count (DL: number of clearances/handover transmissions) were derived from 10 min intervals of more or less constant (maximum) traffic during 30 – 40 min simulation scenarios with different maxima [37]. MWL was measured at the end of 5 min intervals using the workload assessment keypad (WAK, [12]) with a 7-level ATWIT scale (Air Traffic Workload Input Technique: low traffic = 1,….,excessive (= maximum = 7). Figure 5 a), b) in section 3.2 depicts the derived means across three repeated simulation runs with the reported detailed ATWIT-MWL data for Amarillo approach simulation and for the average across the three airports and ATCOs respectively. The LRL-model based estimate of the cognitive Hill-function constant K_c_ = 1/k = exp(-μ/ν) was derived from the logistic regression MWL-parameter estimates (μ, ν) which agreed with the corresponding heuristic sigmoid parameters in [4]. For the cognitive Hill exponent γ = ρ/ν the TL-sensitivity parameter is required. Reliable DLtask-count data were provided only for one of the three airports (Ardmore) and only three average traffic count levels (<NS>) with associated <MWL> levels (“moderate”, “high”, “unmanageable”) and DL-task count values [37] which could be used for LRL-model based estimates of ρ and γ.

#### Experiment 3

Main goal of this research was the investigation of MWL differences in the new work environment of a Remote Tower Center (RTC) for simultaneous control two remote airports (AP). In a realistic human-in-the-loop (HITL)-RTC simulation environment twelve professional tower controllers as participants controlled traffic (factor 1: variable aircraft (AC) count N_S_ (number of AC in control zone)) under three different work conditions (factor 2, including baseline = standard condition, and two-AP control by single ATCO ). Each controller participated in eight repeated simulation runs of 25 min duration under each of the different conditions [50] [2]. Besides a console with standard equipment like flight strips, weather display, control zone radar, and radio communication with (pseudo-) pilots in a separate room, the work environment contained two high resolution video-panoramas, one above the other, which replaced the usual direct view out-of-windows. For the present purpose we used from the whole data set only the data for baseline work condition where the ATCO controlled a single airport (task requirements like in a conventional tower). Like in experiments 1 and 2, besides periodic subjective ISA-MWL reporting (2 min reporting intervals) also communication between ATCOs and pseudo- pilots (count of radio calls per time interval) as main task requirement and call duration was monitored synchronously with ISA-reporting. Traffic count N_S_ as independent environmental simulation variable for each 2 min ISA-interval was derived from the overlapping time intervals of active AC (controlled = monitored callsigns within control zone) between initial call (from pilot entering control zone) until transfer to apron control, resulting in typically 40 to 100 (ISA, RC) data-pairs (N_S_, RC) per traffic count 1 ≤ N_S_ ≤ 10.

### 2. Data

Pre-processed data tables used for the regression analysis in experiments 2 and 3 are provided which are not easily available. Data for experiment 1 are available via open access to ref. [1].

#### Experiment 2: Reconstructed ATWIT-WL Data of [4]

Numerical MWL-data in the following tables from approach-trafffic HITL-simulation of Amarillo, Wichita Falls, Ardmore, (the latter including means across the three airports) were reconstructed from the three data sets depicted in Figure 4 in Lee (2005) in [4]. The discrete aircraft (AC) count N_S_ and ATWIT scale (1…7) for each 5 min simulation interval allowed to extract from Fig. 4 the single MWL-report values per controller (ATCO) for the respective sector: increasing column number = increasing ATWIT-MWL value. Fig. 5a) in section 3.2 depicts the “means”-column listed in the “Amarillo Approach” table. Fig. 5b) depicts the overall means across the three approach sector traffic count means listed in the last column of the “Ardmore Approach” table.

| ATWIT-MWL reports: Amarillo approach sector |  |  |  |  |  |
| --- | --- | --- | --- | --- | --- |
| $N_s$ / peak count per 5 min | MWL-level (4 runs per ATCO) | | | Number of reports | Amarillo Means |
| 5 | NaN | NaN | NaN |  | NaN |
| 6 | NaN | NaN | NaN |  | NaN |
| 7 | 1 | NaN | NaN | 1 | 1,0 |
| 8 | NaN | NaN | NaN |  | NaN |
| 9 | NaN | NaN | NaN |  | NaN |
| 10 | 1 | 2 | NaN | 2 | 1,5 |
| 11 | 1 |  |  | 1 | 1,0 |
| 12 |  |  |  |  | NaN |
| 13 | 1 |  |  | 1 | 1,0 |
| 14 | 2 |  |  | 1 | 2,0 |
| 15 | 1 |  |  | 1 | 1,0 |
| 16 | 2 |  |  | 1 | 2,0 |
| 17 |  |  |  |  | NaN |
| 18 |  |  |  |  | NaN |
| 19 | 3 |  |  | 1 | 3,0 |
| 20 | 3 | 4 |  | 2 | 3,5 |
| 21 | 4 |  |  | 1 | 4,0 |
| 22 | 4 | 5 |  | 2 | 4,5 |
| 23 | 3 | 5 | 6 | 3 | 4,7 |
| 24 | 4 | 5 |  | 2 | 4,5 |
| 25 | 4 |  |  | 1 | 4,0 |
| 26 | 5 |  |  | 1 | 5,0 |
| 27 | 5 |  |  | 1 | 5,0 |
| 28 |  |  |  |  | NaN |
| 29 |  |  |  |  | NaN |
| 30 |  |  |  |  | NaN |
| 31 | 5 |  |  | 1 | 5,0 |

| ATWIT-MWL reports: Wichita Falls approach sector |  |  |  |  |  |  |
| --- | --- | --- | --- | --- | --- | --- |
| n / peak<br>count per 5<br>min | MWL-level (4 runs per ATCO) |  |  |  | Number of<br>reports | Whichia<br>means |
| 5 | NaN | NaN | NaN | NaN |  | NaN |
| 6 | 2 | NaN | NaN | NaN | 1 | 2,0 |
| 7 | NaN | NaN | NaN | NaN |  | NaN |
| 8 | NaN | NaN | NaN | NaN |  | NaN |
| 9 | 2 | NaN | NaN | NaN | 1 | 2,0 |
| 10 | NaN | NaN | NaN | NaN |  | NaN |
| 11 | 1 | 3 | NaN | NaN | 2 | 2,0 |
| 12 |  | NaN | NaN | NaN |  | NaN |
| 13 | 3 | NaN | NaN | NaN | 1 | 3,0 |
| 14 | 3 | 4 | NaN | NaN | 2 | 3,5 |
| 15 | 3 | 4 | 5 | NaN | 3 | 4,0 |
| 16 | 3 | 4 | 5 | 6 | 4 | 4,5 |
| 17 | 4 | 5 | 6 | NaN | 3 | 5,0 |
| 18 | NaN | NaN | NaN | NaN |  | NaN |
| 19 | 5 | 6 | NaN | NaN | 2 | 5,5 |
| 20 | 4 | NaN | NaN | NaN | 1 | 4,0 |
| 21 | 4 | 6 | NaN | NaN | 2 | 5,0 |
| 22 | 6 | NaN | NaN | NaN | 1 | 6,0 |
| 23 | NaN | NaN | NaN | NaN |  |  |
| 24 | NaN | NaN | NaN | NaN |  |  |
| 25 | NaN | NaN | NaN | NaN |  |  |
| 26 | NaN | NaN | NaN | NaN |  |  |
| 27 | NaN | NaN | NaN | NaN |  |  |
| 28 | NaN | NaN | NaN | NaN |  |  |
| 29 | NaN | NaN | NaN | NaN |  |  |
| 30 | NaN | NaN | NaN | NaN |  |  |
| 31 | NaN | NaN | NaN | NaN | NaN | NaN |

| ATWIT-MWL reports: Ardmore approach sector |  |  |  |  |  | Wichita | Amarillo | Means |
| --- | --- | --- | --- | --- | --- | --- | --- | --- |
| n / peak<br>count per 5<br>min | MWL-level<br>(4 runs per ATCO) |  |  | Number<br>of<br>reports | Ardmore<br>WL-Means | Wichita<br>WL-Means | Amarillo<br>Means | across 3<br>sectors<br>(ATCOs) |
| 5 | NaN | NaN | NaN |  | NaN | NaN | NaN | NaN |
| 6 | NaN | NaN | NaN |  | NaN | 2,0 | NaN | 2,0 |
| 7 | 2 | 3 | NaN | 2 | 2,5 | NaN | 1,0 | 1,8 |
| 8 | NaN | NaN | NaN |  | NaN | NaN | NaN | NaN |
| 9 | NaN | NaN | NaN |  | NaN | 2,0 | NaN | 2,0 |
| 10 | NaN | NaN | NaN | 2 | NaN | NaN | 1,5 | 1,5 |
| 11 | 3 | NaN | NaN | 1 | 3,0 | 2,0 | 1,0 | 2,0 |
| 12 | NaN | NaN | NaN |  | NaN | NaN | NaN | NaN |
| 13 | 4 | NaN | NaN | 1 | 4,0 | 3,0 | 1,0 | 2,7 |
| 14 | 2 | 3 | NaN | 2 | 2,5 | 3,5 | 2,0 | 2,7 |
| 15 | 2 | 3 | NaN | 2 | 2,5 | 4,0 | 1,0 | 2,5 |
| 16 | 3 | 5 | NaN | 2 | 4,0 | 4,5 | 2,0 | 3,5 |
| 17 | 4 | 5 | NaN | 2 | 4,5 | 5,0 | NaN | 4,8 |
| 18 | 4 | 5 | 6 | 3 | 5,0 | NaN | NaN | 5,0 |
| 19 | 4 | 5 | NaN | 2 | 4,5 | 5,5 | 3,0 | 4,3 |
| 20 | 4 | 5 | NaN | 2 | 4,5 | 4,0 | 3,5 | 4,0 |
| 21 | 3 | 4 | NaN | 2 | 3,5 | 5,0 | 4,0 | 4,2 |
| 22 | 4 | NaN | NaN | 1 | 4,0 | 6,0 | 4,5 | 4,8 |
| 23 | 4 | NaN | NaN | 1 | 4,0 | NaN | 4,7 | 4,3 |
| 24 | 4 | NaN | NaN | 1 | 4,0 | NaN | 4,5 | 4,3 |
| 25 | 4 | 5 | NaN | 2 | 4,5 | NaN | 4,0 | 4,3 |
| 26 | 4 | 5 | NaN | 2 | 4,5 | NaN | 5,0 | 4,8 |
| 27 | NaN |  | NaN |  | NaN | NaN | 5,0 | 5,0 |
| 28 | NaN |  | NaN |  | NaN | NaN | NaN | NaN |
| 29 | NaN |  | NaN |  | NaN | NaN | NaN | NaN |
| 30 | NaN |  | NaN |  | NaN | NaN | NaN | NaN |
| 31 | NaN |  | NaN |  | NaN | NaN | 5,0 | 5,0 |
| MWL-Means: |  |  |  |  | 3,8 | 4,0 | 3,1 | 3,6 |

#### Experiment 3

Mean values across N(n) measurements of 2 min data acquisition intervals, separated for experimental condition c_0_, c_1_, c_2_, of dependent variables ISA-MWL, RC-TL. Work condition c2 (increased TL due do single ATCO controlling 2 airports (AP)) not used for present analysis. Condition c0 and c1 were combined as “standard condition”, i.e. 2 controllers sharing tasks for two variants of simultaneous 2-AP control.

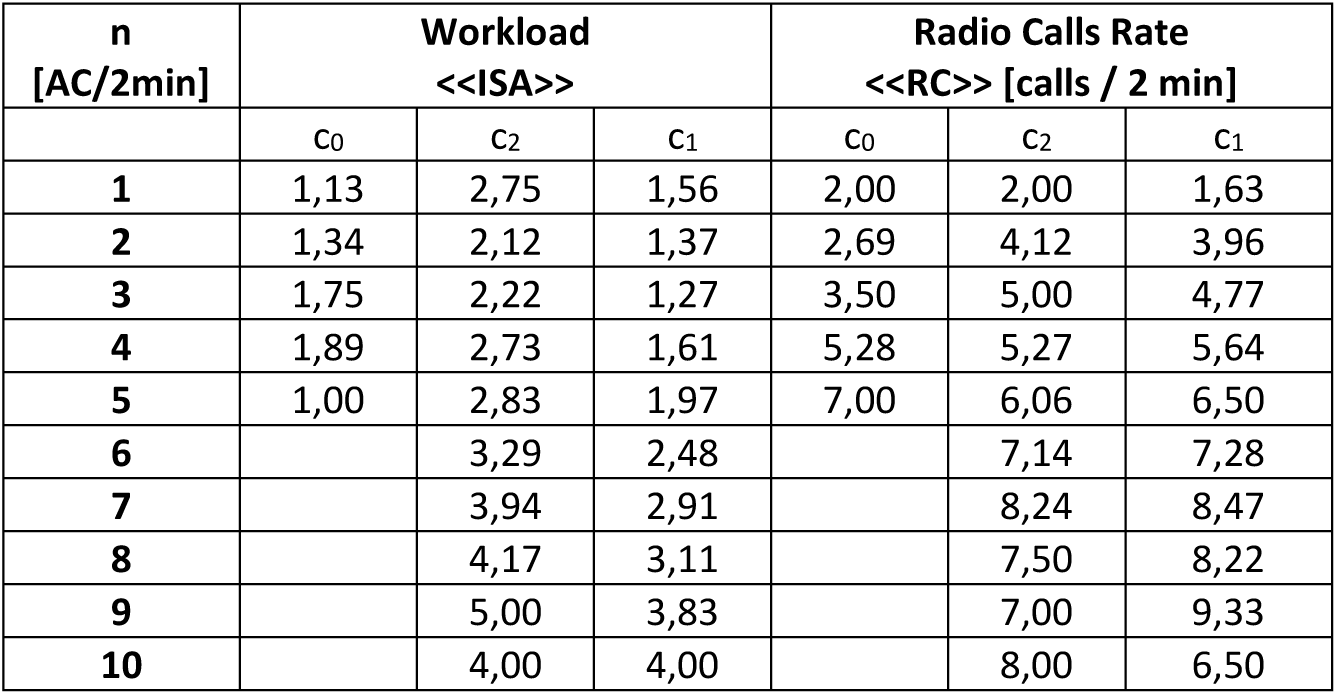

| n<br>[AC/2min] | Workload<br><<ISA>> |  |  | Radio Calls Rate<br><<RC>> [calls / 2 min] |  |  |
| --- | --- | --- | --- | --- | --- | --- |
| | $c_0$ | $c_2$ | $c_1$ | $c_0$ | $c_2$ | $c_1$ |
| 1 | 1,13 | 2,75 | 1,56 | 2,00 | 2,00 | 1,63 |
| 2 | 1,34 | 2,12 | 1,37 | 2,69 | 4,12 | 3,96 |
| 3 | 1,75 | 2,22 | 1,27 | 3,50 | 5,00 | 4,77 |
| 4 | 1,89 | 2,73 | 1,61 | 5,28 | 5,27 | 5,64 |
| 5 | 1,00 | 2,83 | 1,97 | 7,00 | 6,06 | 6,50 |
| 6 |  | 3,29 | 2,48 |  | 7,14 | 7,28 |
| 7 |  | 3,94 | 2,91 |  | 8,24 | 8,47 |
| 8 |  | 4,17 | 3,11 |  | 7,50 | 8,22 |
| 9 |  | 5,00 | 3,83 |  | 7,00 | 9,33 |
| 10 |  | 4,00 | 4,00 |  | 8,00 | 6,50 |

